# Training and match running distances and perceived exertion at sea-level and altitude in elite North American soccer: A natural longitudinal cross-over case series

**DOI:** 10.1101/2025.07.13.664605

**Authors:** Garrison Draper, Paul Chesterton, Tim Thompsion, Matthew D. Wright

## Abstract

**Purpose:** There is limited real-world data on differential perceptions of exertion in professional soccer at altitude. In a natural cross-over design, we quantified responses to training and matches at sea level and altitude.

**Methods:** Total, and high-speed, running distance and differential session ratings of perceived exertion for breathlessness (s-RPE-B) and leg exertion (s-RPE-L) were collected from 19 professional soccer players (age: 27.3 ± 3.4 years, stature: 181.2 ± 5.9cm, mass: 78.7 ± 8.1kg) at sea-level and altitude. Data were collected for matches raining sessions one and two days pre match.

**Results:** Differences between altitude conditions were estimated through mixed-linear modeling, with 95% confidence intervals representing values highly compatible with our observed data. In matches, total distance may have been reduced (95% confidence intervals 43 to -833 m, surprisal [s] = 3.74) and s-RPE-B may have increased (-0.6 to 10.7, s = 3.66) at altitude. A clear perceptual shift was observed with s-RPE-B increasing relative to s-RPE-L (-13.4 to -7.17, s >13) at altitude. Training at altitude included higher total (318 to 661 m, s > 13), and high-speed (33.1 to 93 m, s > 13), running distances but s-RPE-L reduced (-14.2 to -1.01, s = 5.39) and s-RPE-B increased relative to s-RPE-L (-15.1 to -3.93, s = 10.3).

**Conclusions:** Our key finding was breathlessness increased relative to leg exertion at altitude.

## Introduction

Elite North American soccer leagues provide challenging altitude conditions when teams travel to cities such as Salt Lake City, Utah (1356 m above sea level), Denver, Colorado (1609 m), Mexico City, Mexico (2200 m) and Toluca, Mexico (2691 m). Teams based at altitude enjoy significant performance advantages in goals scored and goals conceded over their visiting sea level based opponents. For example, home teams in South America were shown to, on average, have a 0.7 goals per game advantage over the away team when both teams are based at sea-level, however this advantage increases by ∼0.5 goals per game for every 1000 m of altitude the away team travels to (McSharry, 2007). Expert consensus agrees that non-acclimatized players exhibit reduced aerobic performance at these altitudes during soccer games (Bärtsch et al., 2008). A systematic review showed soccer match running performance (total and high-speed distances) may also be reduced although a large heterogeneity of effect sizes have been reported (Draper et al., 2023a). While prior acclimatization may attenuate performance reductions, enabling sufficient time in the environment is often impractical within congested North American soccer schedules (Bärtsch et al., 2008; Draper et al., 2023a).

Maximal oxygen uptake has been consistently shown to be impaired during stays at altitude across multiple studies (Levine et al., 2008). The decline in partial pressure of oxygen (PO_2_) in the atmosphere, initiates a cascade from the alveoli to the arterial blood. This results in arterial oxyhemoglobin saturation which is strongly linked to the ability to maintain maximal oxygen uptake (Chapman, 2013). Exercising at altitude leads to increased blood lactate accumulation and heart rate, as well as subjective measures such as overall perceived exertion and breathlessness (Levine et al., 2008; Young et al., 1982). Maximal oxygen uptake and time to exhaustion in endurance performance appear to decrease linearly from around 500 m of altitude (Cheung, 2010) with a 0.5 to 1% decrease in 𝑉O_₂max_ for every 100 m of simulated altitude (Levine et al., 2008). These effects are influenced by both time spent in the environment and altitude level (Cheung, 2010).

Actions requiring a larger contribution of anaerobic energy pathways are likely unaffected by normobaric hypoxia. Laboratory studies show that single-sprint performance is unaffected by acute exposure to simulated altitude (Girard et al., 2017), despite a reduction in aerobic power (Weyand et al., 1999). It is also hypothesized that reductions in air resistance, based on decreases in air density, may result in improvements in maximal sprinting speed or throwing/kicking for distance at altitude (Girard et al., 2017; Levine et al., 2008). Given that soccer is a repeated, high-intensity, intermittent activity with suboptimal recovery, it is likely any improvements in maximal velocity sprinting may be offset by the increased energetic demand of recovery (Feriche et al., 2007). For example, ATP regeneration is significantly slowed in hypoxia. Altitude (1550 m) was possibly beneficial for rugby players when sprinting 30 m but likely impaired performance in repeated shuttle sprints (Hamlin et al., 2008).

Acute exposures to altitude may affect physical performance outcomes in professional team sport athletes (Draper et al., 2023). Nassis et al. (2010) reported a 3.1% decrease in total distance covered in matches played >1200 m compared to sea level in professional soccer players at the FIFA World Cup in South Africa. Up to a 15% reduction in high-intensity running volumes in high-level soccer players when performing at altitude was observed by Garvican et al. (2014). While these studies highlight external physical demands at altitude, there remains limited data on athletes’ internal physiological responses. Exercising heart rate is elevated upon arrival, with no return to baseline at the completion of a 2-week training camp (Buchheit et al., 2013). From the same case series, Aughey et al. (2013) concluded that changes in physical performance ‘uncoupled’ from match activity at altitude, with reductions in high-intensity capacity markedly more reduced in Australian, compared to acclimatized Bolivian players supporting the notion that physical activity in matches is influenced by contextual and/or tactical factors and not necessarily physical fitness. Therefore, understanding both match running demands and the associated internal physiological responses is crucial. This knowledge helps practitioners optimize preparation strategies and manage performance at altitude.

Ratings of perceived exertion (RPE) are a valid tool for assessing the internal load during soccer training and matches (Impellezzeri et al., 2004; MacPherson et al., 2019; Wright et al., 2020). However, perceived exertion should not be viewed as a gestalt measure and differentiating between central (cardiorespiratory) and peripheral (leg) inputs may provide more specific insight into the perceived internal demands of performing at altitude in professional soccer (Weston et al., 2015; >McLaren et al., 2016a). Aliverti et al. (2011) observed that respiratory RPE tracked respiratory power output during maximal cycling at both sea level and at high altitude (4559 m) although a notable increase was observed at exhaustion. In contrast, while leg RPE at maximal effort was similar between sea level and altitude, cycling power output decreased at altitude, resulting in a relatively higher perceived leg exertion for the same intensity. These authors speculated this was due to a shift in blood flow from the legs to the respiratory muscles (Aliverti et al., 2011). Previous research in military personnel observed significantly higher peripheral (leg), in comparison to breathlessness RPE, in maximal cycling at sea level but not at altitude (4300 m; Young et al., 1982). However, extrapolating cycling studies to running based sports may be problematic given the known differences between central and peripheral exertion between modalities (McLaren et al., 2016). Beyond measuring exercise intensity, RPE may regulate safe and sustainable pacing (Levine and Buono, 2019). It is possible that teams native to altitude have an advantage at altitude because they are better able to regulate pacing throughout the match to avoid excessive or exhaustive exercise (Levine et al., 2008).

Currently, there is limited real-world data on differences in perceptions of exertion in soccer, particularly for teams competing at the highest level of North American soccer who travel to compete at altitude (Draper et al., 2023a). This case series aimed to compare running load and differential RPE during training and matches in elite North American soccer players at sea level and altitude conditions. We hypothesized that perceived cardiorespiratory exertion would increase in altitude conditions.

## Methods

This case series examined the altitude challenges for professional North American soccer players at a single club. The study was approved by Teesside University School of Health and Life Sciences Ethics sub-committee (Study No 238/18) as part of a larger research project. Player data was collected as part of normal club practices between August 2017 and September 2021 and the ethics committee approved secondary data analysis. Players provide written informed consent to the inclusion of anonymized material pertaining to themselves in this research.

### Study Design

Using a natural, single-sample, multiple crossover design athletes completed two to three training sessions and one soccer match within 96 hours of arriving in altitudes ranging from 11 m to 2300 m. Data were collected over a four-year span (2018-2021) during the team’s competitive periods (figure 1).

**Figure 1:**
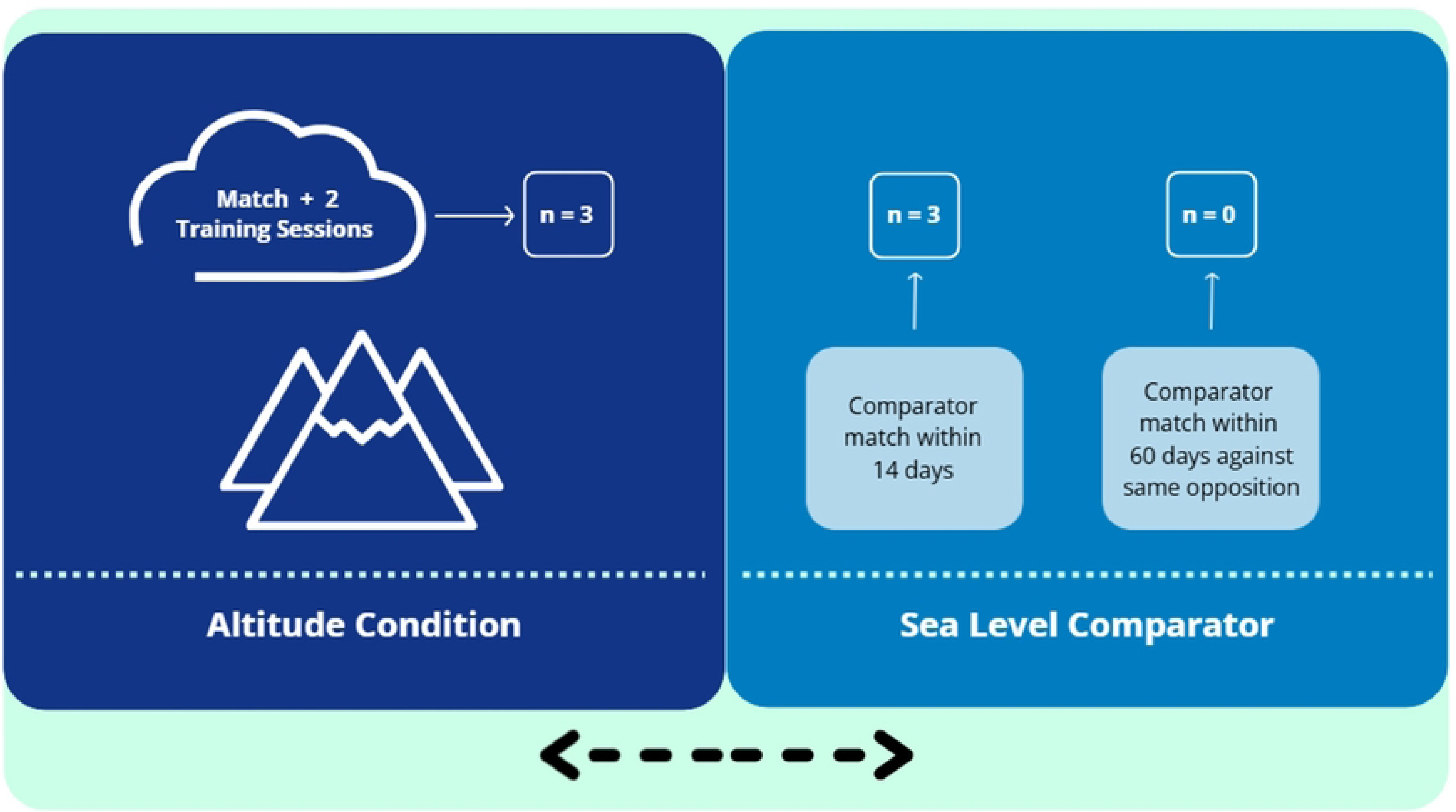
Graphic illustrating the cross-over design with altitude and sea-level comparators.

### Participants

Data from 41 professional soccer players (age: 27.3 ± 3.4 years, stature: 181.2 ± 5.9 cm, body mass: 78.7 ± 8.1 kg), registered at the same professional soccer club competing at the highest level of North American soccer, were included in this study. The roster consisted of players from a range of countries, with approximately 25% playing for their full national teams. Goalkeepers were excluded from the data analyses. Each athlete was required to be available for selection in both the experimental condition (altitude week) and the corresponding control condition (sea-level week) to be considered for data analysis.

### Testing and Data Collection

Data collection occurred during the team’s domestic and continental competitions. The data were split into training day and match day categories, i.e., two days prior to the match (MD-2), one day prior to the match (MD-1), and the day of the match (MD-0). Four selection criteria were set for data inclusion. First, the team needed to play a match at >1000 m of altitude. Second, the altitude match could not fall during a congested fixture period defined as ‘greater than one match per week’ (Bergeron et al., 2012). Third, the subjects needed to complete >59 min on MD-0. Fourth, the control period had to occur within 14 calendar days of the altitude period or be against the same opponent within 60 days of the experimental period. These four criteria resulted in three eligible periods during the four-year window (2018-2021) for analysis. The final data set included three matches at altitude (six training sessions) at altitude and three control matches (six training sessions) which were all against a different opposition and within 14 days of the altitude period. Training sessions lasted a mean duration of 41.2 ± 10.2 minutes.

All data on altitude and environment conditions were collected via the National Weather Service’s website. Altitudes were categorized for analysis based on previous literature (Levine et al., 2008; Alanis et al., 2022), resulting in the following categories: sea level = 0-1000 m, low altitude =1000-2000 m, and moderate altitude = >2000 m. Aerobic performance decrements in athletes have been reported to begin at about 500 m, though most applied literature points to ∼1000 m, up to ∼2000 m (Low Altitude; Bärtsch et al., 2008). Above 2000 m, up to 3000 m (Moderate Altitude) more severe performance decrements can be detected and altitude-based ailments can begin to form, which if left untreated could cause serious medical issues or death (Bergeron et al., 2012). Due to a lack of data points in moderate altitude conditions, we decided to code the data sea level <1000 m and altitude >1000 m.

### Procedures

Global positioning system (GPS) data were collected as part of standard club practices, as described by Draper et al. (2021) via GPS units worn in a harness located between the players’ shoulder blades and sampling at a frequency of 10 Hz, (Optimeye S7, Catapult Innovation, Melbourne, Australia) and total and high-speed running distance (defined as distance covered over 20.2 km/hr) were chosen to represent “volume” and “intensity” (Ishida et al., 2023). Catapult 10-Hz GPS units have demonstrated acceptable validity and good inter-unit reliability [typical error of the measurement 1.3% for total distance and 4.8% for high-speed running] (Scott et al., 2016). Data were only included if it complied with the club’s data standards which checked compliance within the metrics of Horizontal Dilution of Precision (HDOP) (<3) and #Sat (>9) set forth in the Catapult user manual.

Session rates of perceived exertion (s-RPE) were collected following training and games to assess player perceptions of the exertion during the session or game. Players provided ratings representing peripheral lower-limb neuromuscular ‘legs’ exertion (s-RPE-L) and central or cardiorespiratory ‘breathlessness’ exertion (s-RPE-B) using the CR-100 (Borg and Borg, 2002). The CR0-100 scale has been shown to be valid for quantification of training load in elite soccer and may provide more precise measures compared to the CR-10 scale (Fanchini et al., 2016). Breathlessness s-RPE has been shown to be greater in maximal treadmill running exercise, compared to cycling, when heart rate and VO2max are higher, providing a valid measure of central or cardiorespiratory exertion (McLaren et al., 2016). Conversely, leg s-RPE has been shown to be higher in cycling exercise, compared to running, where blood lactate is increased (McLaren et al., 2016) and in resistance exercise compared to intermittent fitness or soccer activity (Wright et al., 2020). Players rated their s-RPE using the CR100 scale with recognized data collection procedures (McLaren et al., 2021) including player education and habituation to the CR-100 scale, the use of a tablet application to reduce risk of conscious bias, and players were always shown the scale. Data were collected within two hours of the session or match using a mobile phone device.

#### Statistical Analysis

Data were imported in R-Studio and a mixed linear model was built to understand the effects of altitude on internal and external training loads. Altitude was set as a fixed factor and coded as sea level, low, or moderate with repeated measures variables nested within the individual player ID. However, several contextual factors also required consideration within the statistical model for the player-reported data. For instance, some players had repeated exposure to the same altitude environments as they had participated in two or three matches at the same opposition venue; these types of data were coded as ‘replications’. In addition, data from training and match days increased the variability in the outcome measures. These data were analyzed separately for each training day (MD-2 and MD-1) and each match day (MD-0).

Analysis involved exploring the effects of accounting for session duration, replication numbers, or both as covariates within the model using the “performance” package. Including session duration in the model significantly improved the model fit across outcome measures (*p* <0.001 to 0.03), except for s-RPE legs (p = 0.53), although this still ranked as the best model. Replication did not improve the model fit (*p* = ∼1.0). The final model included altitude code as a fixed factor, session duration as a covariate and player ID as a random factor. Given altitude may have diverging effects on cardio-respiratory and neuromuscular demands of training and match play, we were not only interested in how these two measures may change at altitude but also change relative to each other (Young et al., 1982), i.e., is there a shift from predominantly neuromuscular to predominantly cardio-respiratory response to matches and training. Here s-RPE-breathlessness was subtracted from s-RPE-legs and coded s-RPE-difference.

Estimated marginal mean differences were calculated using Tukey post hoc Holm’s corrections using the “emmeans” package in R-studio and presented as raw effect sizes with 95% confidence intervals (95% CIs). The values are interpreted within the 95% confidence interval as “highly compatible with our observed data under the background statistical assumptions” of the model (Rafi and Greenland, 2020). Models were visually inspected using the performance package in R-Studio and generally fit the assumptions of linearity and normality of residuals, although there was some evidence of unequal error variance, which were most likely due to the relative lack of repeated observations in the data set. As such, we performed robust analyses, as recommended by Field and Wilcox (2017). Data were presented as mean differences between altitude levels with uncertainty expressed through 95% confidence intervals. We also report p-values to express the probability of observing an effect as large or larger under the background statistical assumptions including the null hypothesis; where true differences between altitude are zero (Greenland et al., 2016). Surprisal values (s-value) were subsequently calculated by taking the negative base-2 logarithm of the p-value (Rafi and Greenland, 2020). These suggest that, assuming all test assumptions are true, the observed data are no more surprising than x consecutive heads when flipping an unbiased coin. A surprisal value of 4.2 would be no more surprising than throwing four consecutive heads and provide four bits of information against the null (zero) hypothesis, and all other model assumptions. Our approach to interpretation of these data was to avoid dichotomizing observations into significant versus non-significant results (Rafi and Greenland, 2020) and the s-value provides a useful statistic to aid interpretation. A summary of the code used is available [here](https://rpubs.com/mdwright07/altitude).

## Results

Raw data are summarized as box plots with individual data points for matches (figures 2 and 3) and training (figures 4 and 5). Adjusted mean values and estimated marginal mean differences between sea-level and altitude are presented in Table 1 for matches and Table 2 for training. We observed changes compatible with our data between 43 and –818 meters in match total distance at altitude and between -0.6 to 10.7 change in breathlessness exertion (42 observations from 20 players). These represent just under four bits of information (s-values 3.74 and 3.66 respectively) against our model assumptions and no more surprising than three consecutive heads on a fair coin. Changes in high-speed running and s-RPE legs were no more surprising than throwing two consecutive heads (36 observations from 18 players). However, clear differences were observed in the difference between s-RPE breathlessness and s-RPE legs, with breathlessness s-RPE increasing relative to leg s-RPE as athletes moved from sea level to altitude-based competition. Changes between 7.17 and 13.4 Arbitrary Units (AU) were highly compatible with our data.

**Figure 2:**
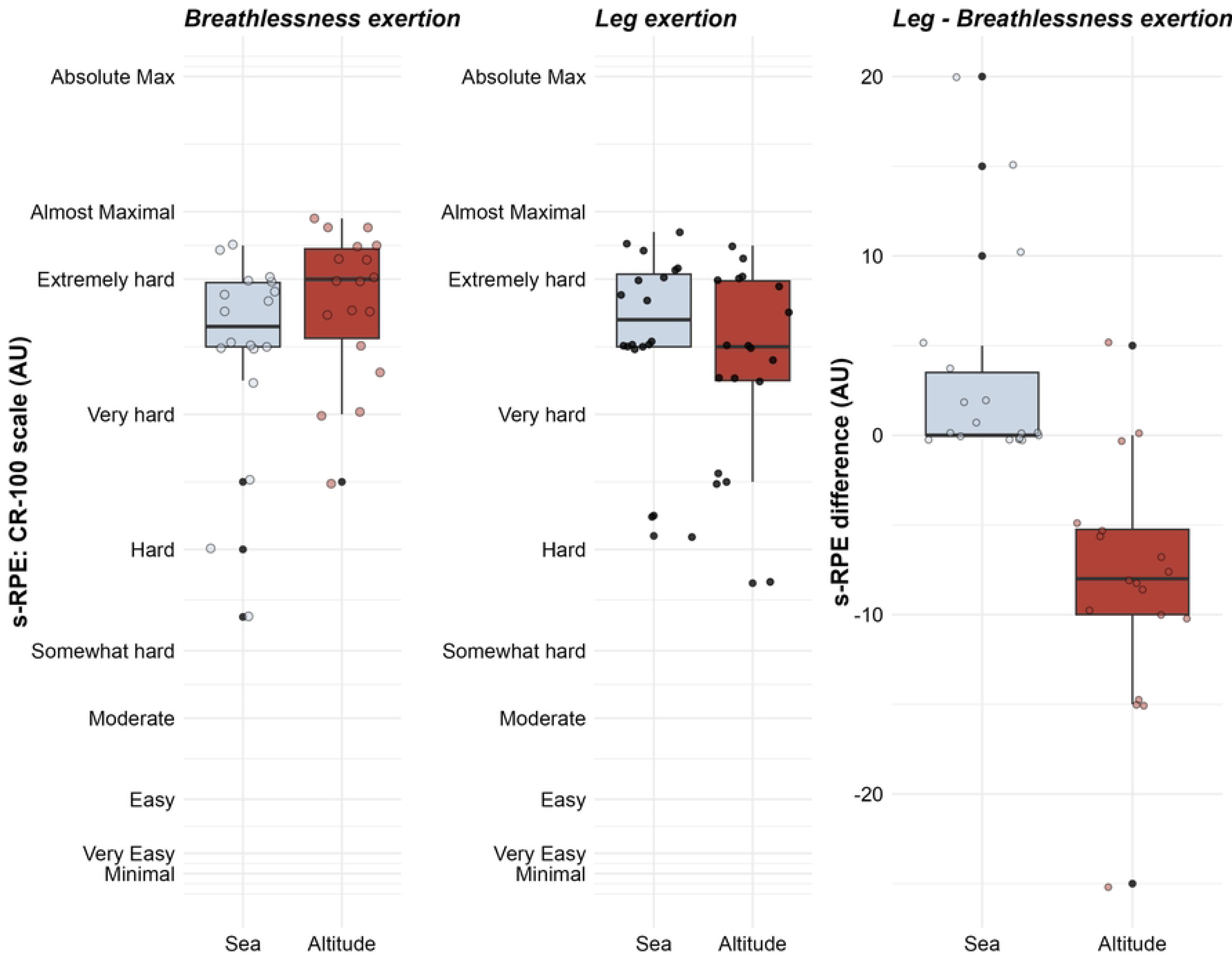
Box and whisker plots visualizing the median values and interquartile range for match s-RPE at sea level and at altitude. Light grey points represent jittered individual data points and black points represent outliers.

**Figure 3:**
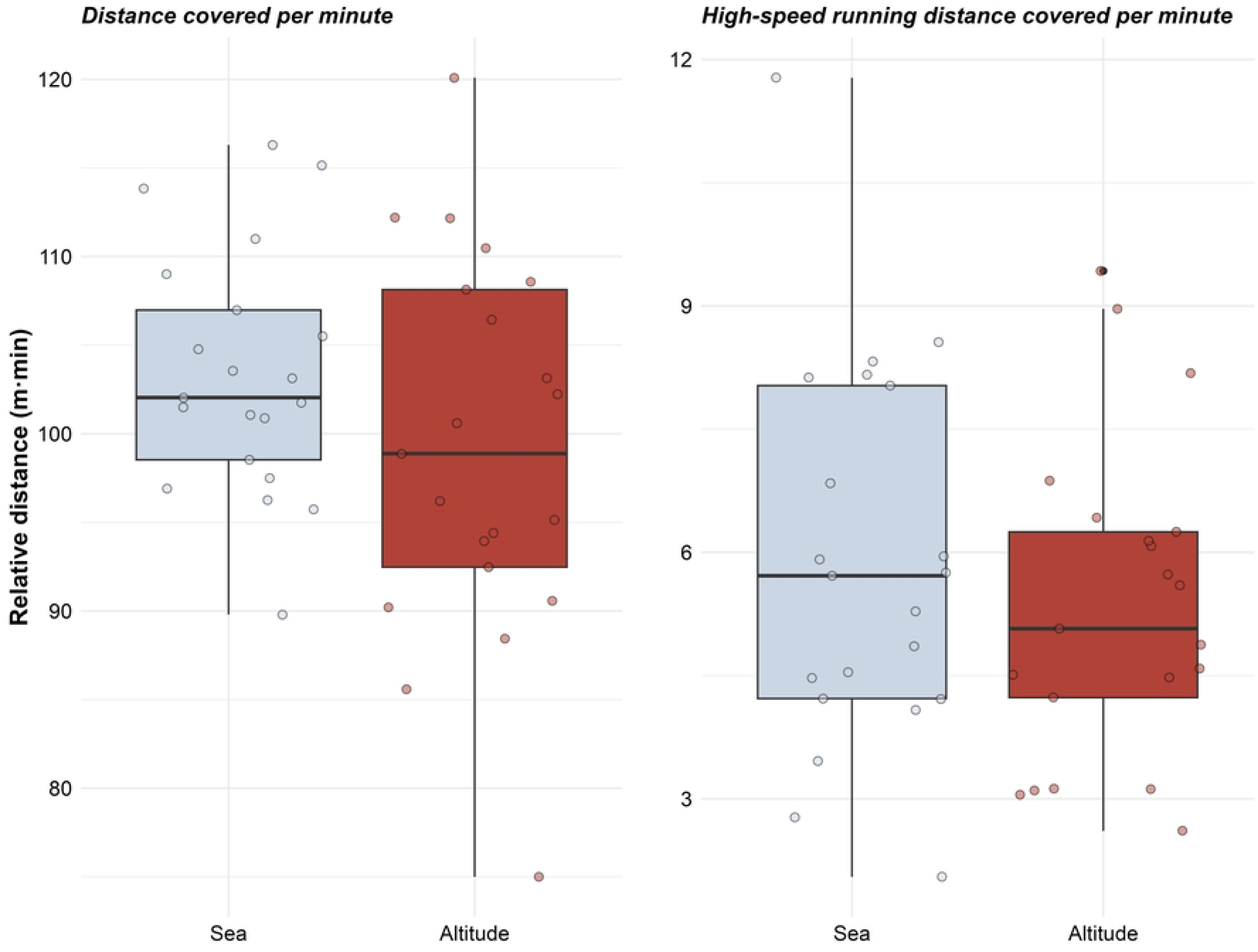
Box and whisker plots visualizing the median values and interquartile range for match running distance at sea level and at altitude. Light grey points represent jittered individual data points and black points represent outliers.

**Table 1:**
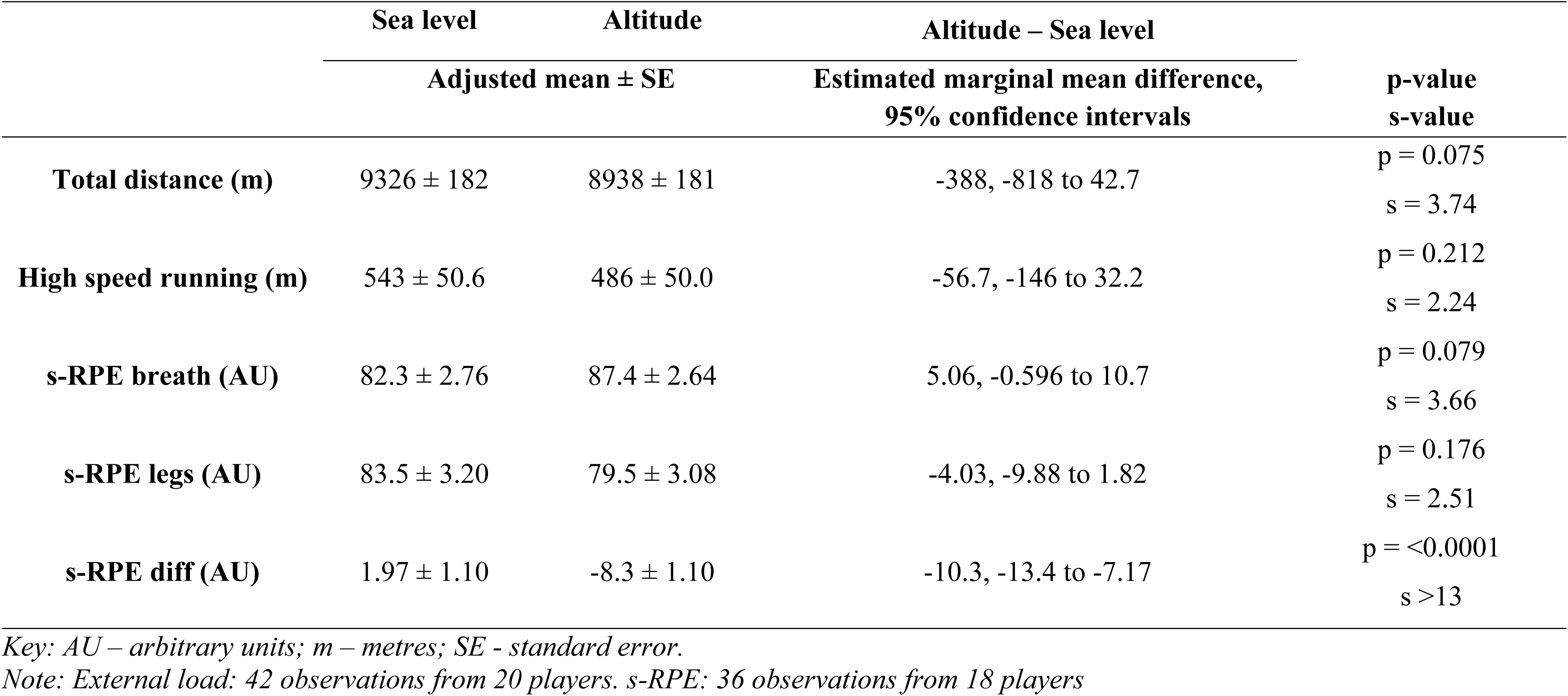
Estimated marginal means for the observed differences between altitudes in matches. The model included duration as a co-variate to account for between- and within-player variation in match minutes.

**Table 2:** Estimated marginal means for the observed differences between altitudes in training. The model included duration as a co-variate to account for between- and within-player variation in match minutes.

|  | Sea level | Altitude | Altitude – Sea level |  |
| --- | --- | --- | --- | --- |
| | Adjusted mean ± SE | | Estimated marginal mean difference,<br>95% confidence intervals | p-value<br>s-value |
| **Total distance (m)** | 2538 ± 63.5 | 3027 ± 65.7 | 490, 318 to 661 | p = <0.0001<br>s = >13 |
| **High speed running (m)** | 77.7 ± 9.64 | 141 ± 10.2 | 63.0, 33.1 to 93.0 | p = <0.0001<br>s = >13 |
| **s-RPE breath (AU)** | 51.2 ± 2.19 | 54.7.4 ± 2.01 | 3.47, -2.80 to 9.74 | p = 0.278<br>s = 1.85 |
| **s-RPE legs (AU)** | 53.4 ± 2.31 | 45.8 ± 1.12 | -7.62, -14.2 to -1.01 | p = 0.024<br>s = 5.39 |
| **s-RPE diff (AU)** | 1.64 ± 1.94 | -7.86 ± 1.78 | 9.50, -15.1 to -3.93 | p = 0.0008<br>s = 10.3 |

*Key: AU – arbitrary units ; m – metres; SE - standard error.*
*Note: External load: 84 observations from 36 players. s-RPE: 67 observations from 32 players*

In training, we observed increases in running of between 318 and 661 m total, and between 33 and 93 m in high-speed running distance, which were highly compatible with our data. Despite this increase in running load, s-RPE legs appeared to be reduced (-14.2 to -1.01 AU, compatible with our data) resulting in a subsequent shift in perceptual response, similar to that observed in matches.

**Figure 4:**
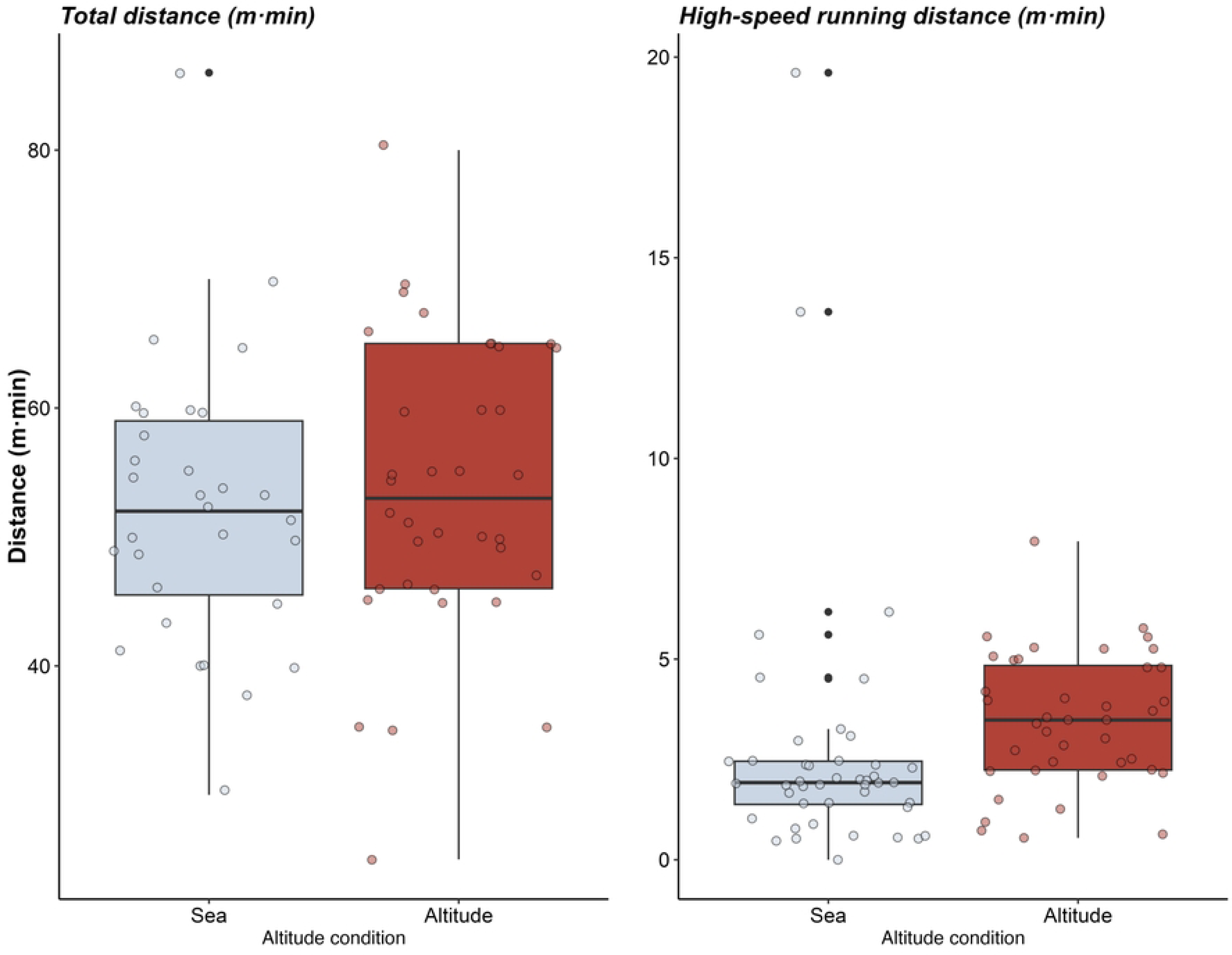
Box and whisker plots visualizing the median values and interquartile range for training s-RPE at sea level and at altitude. Light grey points represent jittered individual data points and black points represent outliers.

**Figure 5:**
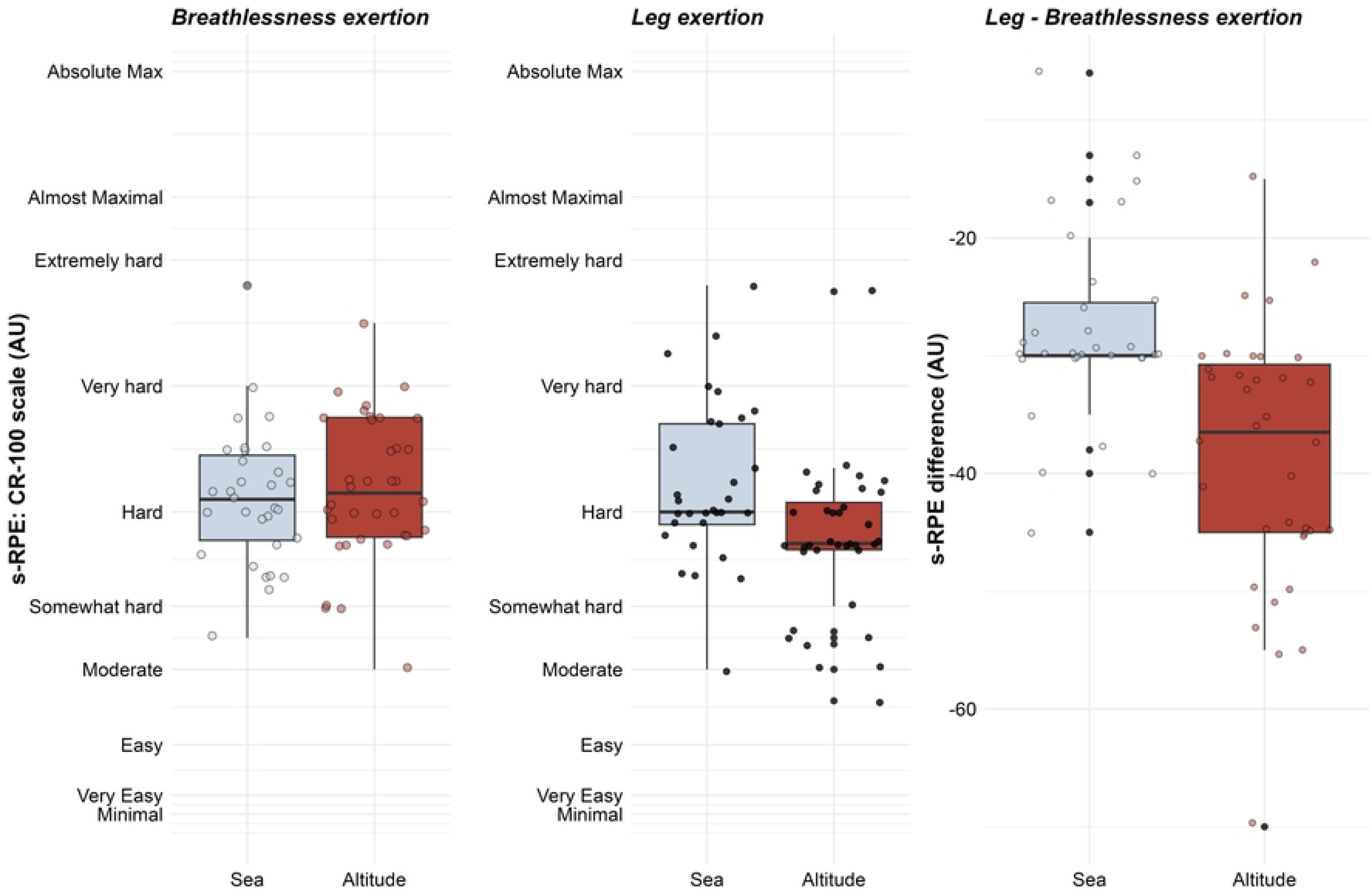
Box and whisker plots visualizing the median values and interquartile range for training running distance at sea level and at altitude. Light grey points represent jittered individual data points and black points represent outliers.

## Discussion

This case series aimed to compare perceived exertion and running training and match load at varying altitudes. We hypothesized that altitude would increase perceptual responses, particularly breathlessness exertion and reduce match running performance. No clear evidence of a change in match running performance was observed but breathlessness s-RPE increased in comparison with leg s-RPE at altitude. In training, players covered more total and high-speed distance at altitude however, leg exertion was reduced both independently and in relation to breathlessness exertion. Our key finding was breathlessness exertion increased relative to leg exertion, suggesting a relative increase in perceived cardiorespiratory (heart rate, oxygen uptake) compared to perceived lower limb, peripheral exertion (neuromuscular or musculoskeletal (McLaren et al., 2016).

In the current dataset, sea level-acclimated athletes played matches at altitude within three to four days of arrival. This arrival fell well outside the recommended time frame for acclimatization, typically referenced as 14-21 days for optimizing performance at altitude (Levine et al., 2008). It would be expected that sea level-acclimated athletes would be susceptible to altitude related changes in performance. For instance, Buchheit et al. (2013) monitored Australian and Bolivian soccer players during a 2-week altitude camp. Australian players, with no altitude experience, showed greater declines in wellness, s-RPE, and yo-yo test performance compared to Bolivian players. Australian players also acclimatized more slowly, suggesting that previous altitude exposure may attenuate negative responses. Although this provides valuable real-world data, it does not account for performance during competitive matches. Draper et al. (2023a) reported in their systematic review of team sport performance at altitude that some measures of physical performance, such as total and high-speed running distances may be impacted by altitude conditions, though the magnitude and duration of the stressors are key determinants of the outcome. We aimed to categorize altitude into low and moderate levels, rather than dichotomizing. Due to a small number of observations at moderate altitudes, all of which were within the typical range of low-altitude values (supplementary figures 1 and 2), justified comparison between altitude and sea level.

The key finding in this study was the perceptual shift in breathlessness and leg exertion. While we did not directly measure aerobic or cardiorespiratory markers (heart rate, O^2^ etc.) breathlessness s-RPE was chosen because it is able to differentiate between exercise with differing cardiorespiratory demands (McLaren et al., 2016). As such our results generally support the strong body of evidence consistently indicating increased cardiorespiratory load when performing at altitude (Levine et al., 2008; Cheung, 2010; Gregson and Drust, 2012), although increases in match breathlessness s-RPE were not clear (s = 3.66 for matches). Altitude exposure also leads to increased blood lactate accumulation, higher heart rates, and greater subjective perceptions of exertion, including breathlessness (Buchheit et al., 2013; Levine et al., 2008; Young et al., 1982). Previous research in military personnel has suggested that differences between leg and breathlessness s-RPE may change at altitude (Young et al., 1982). Physiological measures, such as maximal oxygen consumption, are significantly impaired almost immediately at altitude (Levine et al., 2008). At altitude, lab-based hypoxic cycling results in higher ventilation rates, heart rate, and blood lactate compared to sea level, with a 1% decrease in 𝑉O_₂max_ for every 100 m increase in altitude above 1500 m (Levine et al., 2008).

The extent to which our “real world” data confirms or contradicts these studies is not clear. While breathlessness s-RPE increased in relation to leg s-RPE in both matches and training, a reduction in leg exertion rather than an increase in breathlessness may have contributed to this shift. If our model’s assumptions hold true, we would expect the observed increase in breathlessness s-RPE to be more surprising than the decrease in leg s-RPE (s = 3.66 vs s = 2.51). During soccer training, there was a clear reduction in leg s-RPE despite an increase in total and high-speed running distance and no clear increases in breathlessness s-RPE. This may reflect the lower intensity of training compared to matches or the reduced air resistance due to lower air density at altitude (Levine et al., 2008). Our data aligns with that of Young et al. (1982) who observed local (leg) s-RPE to decrease during maximal cycling at high altitude. Although this somewhat contradicts Aliverti et al. (2011) who report similar leg s-RPE values (∼8 on the CR-10 scale) at volitional exhaustion on an exercise bike, but at a substantially reduced power output (∼50 W) compared to sea level. It is important to note that the altitude conditions in these studies were much higher (over 4300 m) than in our study. This perceptual shift is a novel finding in soccer and warrants further exploration, ideally in larger cohorts across multiple clubs. These data also support the use of differential ratings of perceived exertion, building on previous work showing consistent, modally dependent responses to a training stimulus (McLaren et al., 2017; McLaren et al., 2016; Wright et al., 2020).

Despite taking a quasi-experimental, longitudinal, approach, the reported study has limitations. A key limitation was the number of observations that met our inclusion criteria, which should be considered when interpreting the findings, especially given the natural variability of the outcome measures. As such, we decided to simplify our comparisons to sea level versus altitude and may have lost important nuance with regard to exploring effects across low and moderate altitude zones (Alanis et al., 2022; Bergeron et al., 2012). Another limitation was that the opposition team was not the same in matches in the altitude and sea level condition. We had extended the window in which a sea level comparison period could be included to 60 days (from 14) if the opposition was the same, but unfortunately, there were no such instances. This is an unfortunate limitation, but one that was unavoidable. However, these data suggest some interesting findings that could inform larger studies across multiple teams. Our design could potentially have enabled replication of altitude and sea level conditions in individual participants. This could be important because performance is variable and data on repeated or replicated exposures to altitude could enable the evaluation of individual differences in response to be explored (Chesterton et al., 2021; Senn, 2016). Unfortunately, due to the limited conditions, available matches and the always-changing landscape of professional soccer rosters, there were only a small number of athletes who completed matches in multiple altitude conditions (see supplementary material).

We chose a traditional frequentist approach for statistical analysis, acknowledging the limitations of null-hypothesis significance testing, and aimed to avoid dichotomizing findings into significant versus non-significant results (Rafi and Greenland, 2020). Consideration should be given to the inherently large match-to-match variation in soccer (Gregson et al., 2010). Analysis of previously published data (Draper et al., 2023b) suggested a within player variation in total and high-speed running of 551m (95% confidence intervals, 522-584) and 144m (136 to 152). Despite the limited observations, we found p-values <0.001, which in a crossover study suggests that the sample size was adequate to account for within-player variability. The authors recognize statistical significance does not necessarily denote practical significance, which refers to an effect being large enough to be practically meaningful.

Determining the practically meaningful difference in match-running performance is difficult, as there is no obvious anchor to suggest a minimum threshold associated with a change in match result or physiological markers. In these cases, researchers are often required to default to a distribution approach (0.2 x between-player *SD,* Hopkins et al., 2009) to estimate the smallest worthwhile change (SWC); i.e., 102 m for total distance and 29 m for high-speed running. There are noted shortcomings with the use of SWC, though in the absence of other options, it was determined that this approach was appropriate (Datson et al., 2022). In our study, the mean changes in running performance exceeded these thresholds, suggesting that statistically significant results were also practically meaningful. A practically relevant change in s-RPE of 8 AU (arbitrary units) on the CR100 scale has been established (Wright et al., 2020). This is the average minimal threshold that would be associated with a change in verbal anchor on the scale, i.e., from “somewhat hard” to “hard”. Practically meaningful changes in s-RPE difference (± 8 AU), are compatible with our data in both training and matches.

### Practical Applications

This study demonstrated a shift in athletes’ perceptions of exertion, increasing breathlessness relative to leg exertion when playing and training during acute visits to altitude. Peripheral exertion experienced in training appears to reduce at altitude even when running distances increase. Practitioners working with teams, who travel to altitude should consider differentiating s-RPE to better monitor specific exertion responses. These data may help practitioners in the preparation of athletes to compete at altitude with limited acclimatization time available, as well as subsequent recovery and nutrition needs.

## Conclusion

This study aimed to evaluate match and training loads at both sea level and altitude. Breathlessness s-RPE increased relative to leg exertion in both training and matches when running distances were not clearly different. Running demands of training were increased at altitude, but this was accompanied by a decrease in leg exertion. The methodological approach, including the collection of high-quality differential s-RPE data (McLaren et al., 2021), could be replicated in future research across multiple teams.

## Declaration of Funding

No funding was received for this project

## Figure legends

Supplementary figure 1: Box and whisker plots visualizing the median values and interquartile range for match s-RPE at sea level, low and moderate altitude. Light grey points represent jittered individual data points and black points represent outliers.

Supplementary figure 2: Box and whisker plots visualizing the median values and interquartile range for match running distance at sea level, low and moderate altitude. Light grey points represent jittered individual data points and black points represent outliers.

## Notes

### Competing Interest Statement

The authors have declared no competing interest.

## REFERENCES

Alanis A, Salas O, Salas K, Quintero I, Carranza Y, Salazar L. Playing at altitude. Performance of a Mexican professional football team at different level of altitude. Apunts Sports Med, 2022, 1;57(215):100391.

Aliverti, A., Kayser, B., Mauro, A.L., Quaranta, M., Pompilio, P., Dellacà, R.L., Ora, J., Biasco, L., Cavalleri, L., Pomidori, L. and Cogo, A., 2011. Respiratory and leg muscles perceived exertion during exercise at altitude. Respiratory physiology & neurobiology, 177(2), 162 – 168.

Aughey RJ, Hammond K, Varley MC, Schmidt WF, Bourdon PC, Buchheit M, Simpson B, Garvican-Lewis LA, Kley M, Soria R, Sargent C. Soccer activity profile of altitude versus sea-level natives during acclimatisation to 3600 m (ISA3600). Br J Sports Med. 2013;47(Suppl 1):i107–13.

Bärtsch P, Saltin B, Dvorak J. Consensus statement on playing football at different altitude. Scand J Med Sci Sports, 2008;18(Suppl.1):96–99.

Bergeron MF, Bahr R, Bärtsch P, Bourdon L, Calbet JA, Carlsen KH, Castagna O, González-Alonso J, Lundby C, Maughan RJ, Millet G. International Olympic Committee consensus statement on thermoregulatory and altitude challenges for high-level athletes. Br J Sports Med. 2012;46(11):770–9.

Borg E, Borg G. A comparison of AME and CR100 for scaling perceived exertion. Acta psychologica. 2002 Feb 1;109(2):157–75.

Buchheit M, Simpson BM, Garvican-Lewis LA, Hammond K, Kley M, Schmidt WF, Aughey RJ, Soria R, Sargent C, Roach GD, Claros JC. Wellness, fatigue and physical performance acclimatisation to a 2-week soccer camp at 3600 m (ISA3600). Br J Sports Med. 2013;47(Suppl 1):i100–6.

Chapman RF. The individual response to training and competition at altitude. Br J Sports Med. 2013;47(Suppl 1):i40–4.

Chesterton P, Evans W, Wright M, Lolli L, Richardson M, Atkinson G. Influence of lumbar mobilizations during the Nordic hamstring exercise on hamstring measures of knee flexor strength, failure point, and muscle activity: a randomized crossover trial. J Manip Physiol Ther. 2021;44(1):1–3.

Cheung S. Advanced Environmental Exercise Physiology. Edited by M. Zavala. Champaign, IL: Human Kinetics; 2010.

Datson N, Lolli L, Drust B, Atkinson G, Weston M, Gregson W. Inter-methodological quantification of the target change for performance test outcomes relevant to elite female soccer players. Sci Med Football. 2022;6(2):248–61.

Draper G, Atkinson G, Chesterton P, Portas M, Wright M. Elite North American soccer performance in thermally challenging environments: An explorative approach to tracking outcomes. J Sports Sci. 2023b;41(11):1107–14.

Draper G, Wright MD, Ishida A, Chesterton P, Portas M, Atkinson G. Do environmental temperatures and altitudes affect physical outputs of elite football athletes in match conditions? A systematic review of the ‘real world’ studies. Sci Med Football. 2023a;7(1):81–92.

Draper G, Wright, M, Chesterton P, Atkinson G. The tracking of internal and external training loads with next-day player-reported fatigue at different times of the season in elite soccer players. Int J Sports Sci Coach, 2021;16:793–803

Feriche B, Delgado M, Calderón C, Lisbona O, Chirosa IJ, Miranda MT, Fernandez JM, Alvarez J. The effect of acute moderate hypoxia on accumulated oxygen deficit during intermittent exercise in nonacclimatized men. J Strength Cond Res. 2007;21(2):413–8.

Field AP, Wilcox RR. Robust statistical methods: A primer for clinical psychology and experimental psychopathology researchers. Behav Res Ther. 2017;98:19–38.

Fanchini, M., Ferraresi, I., Modena, R., Schena, F., Coutts, A. J., & Impellizzeri, F. M. Use of the CR100 scale for session rating of perceived exertion in soccer and its interchangeability with the CR10. Int J Sports Physiol Perf. 2016; 11(3), 388–392.

Garvican LA, Hammond K, Varley MC, Gore CJ, Billaut F, Aughey RJ. Lower running performance and exacerbated fatigue in soccer played at 1600 m. Int J Sports Physiol Perf. 2014;9(3):397–404.

Girard, O., Brocherie, F. and Millet, G.P., 2017. Effects of altitude/hypoxia on single-and multiple-sprint performance: a comprehensive review. Sports medicine, 47, pp.1931–1949.

Greenland S, Senn SJ, Rothman KJ, Carlin JB, Poole C, Goodman SN, Altman DG. Statistical tests, P values, confidence intervals, and power: a guide to misinterpretations. Eur. J. Epidemiol. 2016;31(4):337–50.

Gregson W, Drust B, Atkinson G, Salvo VD. Match-to-match variability of high-speed activities in premier league soccer. Int J Sports Med. 2010:237–42.

Gregson, W. and Drust, B. Environmental stress, In Williams MA, Ford P, Drust B. Science and Soccer: Developing Elite Performers, Third Edition. London, Routledge; 2012

Hamlin, M.J., Hinckson, E.A., Wood, M.R. and Hopkins, W.G., 2008. Simulated rugby performance at 1550-m altitude following adaptation to intermittent normobaric hypoxia. Journal of Science and Medicine in Sport, 11(6), pp.593–599.

Hopkins W, Marshall S, Batterham A, Hanin J. Progressive statistics for studies in sports medicine and exercise science. Med Sci Sport Exerc. 2009;41(1):3.

Impellizzeri, F.M., Rampinini, E., Coutts, A.J., Sassi, A.L.D.O. and Marcora, S.M., 2004. Use of RPE-based training load in soccer. Medicine & Science in sports & exercise, 36(6), pp.1042–1047.

Ishida A, Draper G, Wright M, Emerson J, Stone MH. Training Volume and High-Speed Loads Vary Within Microcycle in Elite North American Soccer Players. J Strength Cond Res. 2023;37(11):2229–34.

Levine BD, Stray-Gundersen J, Mehta RD. Effect of altitude on football performance. Scand J Med Sci Sports, 2008;18(Suppl.1):76–84.

Levine, A. and Buono, M.J., 2019. Rating of perceived exertion increases synergistically during prolonged exercise in a combined heat and hypoxic environment. Journal of thermal biology, 84, pp.99–102.

Macpherson, T.W., McLaren, S.J., Gregson, W., Lolli, L., Drust, B. and Weston, M., 2019. Using differential ratings of perceived exertion to assess agreement between coach and player perceptions of soccer training intensity: an exploratory investigation. Journal of sports sciences, 37(24), pp.2783–2788.

McLaren, SJ., Coutts, A. J., & Impellizzeri, F. M. (2021). Perception of effort and subjective monitoring. In French D, Ronda LT, editors. NSCA’s Essentials of Sport Science. Champaign, IL: Human Kinetics; 2021

McLaren, S.J., Graham, M., Spears, I.R. and Weston, M., 2016. The sensitivity of differential ratings of perceived exertion as measures of internal load. International Journal of Sports Physiology and Performance, 11(3), pp.404–406.

McLaren, S.J., Smith, A., Spears, I.R. and Weston, M., 2017. A detailed quantification of differential ratings of perceived exertion during team-sport training. Journal of science and medicine in sport, 20(3), pp.290–295.

McSharry PE. Altitude and athletic performance: statistical analysis using football results. Br Med J, 2007;335(7633):1278-81.

Nassis GP. Effect of altitude on football performance: analysis of the 2010 FIFA World Cup Data. J Strength Cond Res. 2013;27(3):703–7.

Rafi Z, Greenland S. Semantic and cognitive tools to aid statistical science: replace confidence and significance by compatibility and surprise. BMC Med Res Methodol. 2020;20:1–3.

Scott MTU, Scott TJ, Kelly VG. The validity and reliability of global positioning systems in team sport: A brief review. J Strength Cond Res 30: 1470–1490, 2016

Senn S. Mastering variation: variance components and personalised medicine. Stat Med. 2016;35(7):966–77.

Weston, M., Siegler, J., Bahnert, A., McBrien, J. and Lovell, R., 2015. The application of differential ratings of perceived exertion to Australian Football League matches. Journal of science and medicine in sport, 18(6), pp.704–708.

Wright MD, Songane F, Emmonds S, Chesterton P, Weston M, McLaren SJ. Differential ratings of perceived match and training exertion in girls’ soccer. Int J Sports Physiol Perform. 2020;15(9):1315–23.

Young AJ, Cymerman A, Pandolf KB. Differentiated ratings of perceived exertion are influenced by high altitude exposure. Med Sci Sports & Exer. 1982;14(3):223–8.

